# Horizontal transferred T-DNA and haplotype-based phylogenetic analysis uncovers the origin of sweetpotato

**DOI:** 10.1101/2022.09.30.510208

**Authors:** Mengxiao Yan, Ming Li, Yunze Wang, Xinyi Wang, M-Hossein Moeinzadeh, Dora G. Quispe-Huamanquispe, Weijuan Fan, Yuqin Wang, Haozhen Nie, Zhangying Wang, Bettina Heider, Robert Jarret, Jan F. Kreuze, Godelieve Gheysen, Hongxia Wang, Ralph Bock, Martin Vingron, Jun Yang

## Abstract

The hexaploid sweetpotato is one of the most important root crops worldwide. However, its genetic origins are controversial. In this study, we identified two progenitors of sweetpotato by horizontal gene transferred *Ib*T-DNA and haplotype-based phylogenetic analysis. The diploid progenitor is the diploid form of *I. aequatoriensis*, contributed the B_1_ subgenome, *Ib*T-DNA2 and lineage 2 type of chloroplast genome to sweetpotato. The tetraploid progenitor of sweetpotato is *I. batatas* 4x, donating the B_2_ subgenome, *Ib*T-DNA1 and lineage 1 type of chloroplast genome. Sweetpotato derived from the reciprocal cross between the diploid and tetraploid progenitors and a subsequent whole genome duplication. We also detected biased gene exchanges between subgenomes. The B_1_ to B_2_ subgenome conversions were almost 3-fold higher than the B_2_ to B_1_ subgenome conversions. This study sheds lights on the evolution of sweetpotato and paves a way for the improvement of sweetpotato.

## Introduction

Sweetpotato, *Ipomoea batatas* (L.) Lam. (2n = 6x = 90), was first domesticated in tropical America at least 5000 years ago ^1^, introduced into Europe and Africa in the early 16^th^ century, and later into the rest of the world ^2^. Today, sweetpotato has become an important staple crop worldwide with an annual production of ∼113 million tons, and is an important source of dietary calories, proteins, vitamins and minerals. For example, orange-fleshed sweetpotato plays a crucial role in combating vitamin A deficiency in Africa ^3–5^. Unlike other important polyploid crops, such as hexaploid bread wheat (*Triticum aestivum*) and tetraploid potato (*Solanum tuberosum*), the origin of cultivated sweetpotato has been the subject of considerable debate. Furthermore, the exact role of polyploidization in the origin and evolution of sweetpotato has not been determined. Knowledge of its genetic origin is vital for supporting further studies of its biology, domestication, genetics, genetic engineering and breeding using of its wild relatives.

Sweetpotato belongs to the genus *Ipomoea* series Batatas (Convolvulaceae). This group includes *I. batatas* 6x (sweetpotato), *I. batatas* 4x, and 13 diploid species that are commonly considered as the wild relatives of the cultivated sweetpotato ^6^. Three polyploidization scenarios have been proposed to account for the hexaploid genome of sweetpotato. First, the autopolyploid hypothesis suggests that sweetpotato has an autopolyploid origin with *I. trifida* being the only progenitor. This hypothesis has gained supports from phylogenetic analysis ^7,8^, polysomic inheritance based on genetic linkage analysis ^9–13^, and cytogenetic analyses ^14,15^. Second, Gao, et al. ^16^ postulated that the hexaploid sweetpotato may be a segmental allopolyploid based on the analysis of 811 conserved single-copy genes. Third, the allopolyploidy hypotheses are diverse and relatively less consistent. Based on cytogenetic analysis, Nishiyama ^17^ suggested that sweetpotato originated from *I. trifida* 3x, which is a hybrid between *I. leucantha* and *I. littoralis*. Austin ^1^ suggested that the cultivated sweetpotato was derived from a hybridization event between *I. trifida* and *I. triloba* based on morphological data. Gao et al. ^18^, based on *Waxy* (*Wx*) intron sequence variation, suggested that sweetpotato arose via hybridization between *I. tenuissima* and *I. littoralis*. However, both cytogenetic and recent genomic analyses suggest that sweetpotato (B_1_B_1_B_2_B_2_B_2_B_2_) composed of two subgenomes and arose from a cross between a diploid and a tetraploid progenitor ^19,20^. The diploid progenitor is most likely *I. trifida*, whereas the tetraploid progenitor has remained debated ^21^. Based on a phylogenetic analyses of homologous haplotypes, Yan et al. ^22^ suggested the tetraploid progenitor of sweetpotato is *I. batatas* 4x. However, Muñoz-Rodríguez et al. ^23^ identified *I. aequatoriensis* as the tetraploid progenitor of sweetpotato based on morphological and phylogenetic analyses. Therefore, the origin of sweetpotato is still controversial and need to be determined.

As the first reported natural transgenic food crop, the genomes of almost all sweetpotato cultivars/landraces and some of its wild relatives contain horizontally transferred *Ib*T-DNA1 and/or *Ib*T-DNA2 sequences from *Agrobacterium* spp. ^24,25^. *Ib*T-DNAs were inherited from the progenitors and could serve as natural genetic markers to track the progenitors of cultivated sweetpotato ^24^. Therefore, the *Ib*T-DNA positive species in the series Batatas (*I. trifida, I. cordatotriloba, I. tenuissima*, and *I. batatas* 4x) are potential wild progenitors of sweetpotato ^24^. Consequently, in order to trace the genetic origin(s) of cultivated sweetpotato, *I. batatas* 4x and other wild relatives in the *Ipomoea* series Batatas are key species to be examined ^24^.

Because of the highly heterozygous and complex hexaploid genome ^5,26^, a serious limitation in most previous studies on the genetic origin of sweetpotato has been the use of consensus genomic sequences and a limited number of nuclear markers. In addition, chromosome rearrangements and homoeologous exchanges that shuffle and/or replace homoeologs among the subgenomes of polyploids ^27–29^ further complicate genetic studies that aim to resolve the origin of polyploid species. Currently, the best strategy for determining the origin of allopolyploids relies on the use of subgenome-level genome assemblies or the the homologous genes or variants of each subgenome to perform the phylogenetic analyses. This strategy has been successfully applied to rapeseed (*Brassica napus*), bread wheat (*Triticum aestivum*) and *Echinochloa* spp., polyploid bamboo (*Bambusa* spp.) and strawberry (*Fragaria × ananassa*) ^30–35^. However, unlike these allopolyploids, the subgenomes of sweetpotato are highly similar to one another due to the close genetic relationship between the diploid and the tetraploid progenitor species. Also, a subgenome-level or fully-phased reference genome of sweetpotato is not yet available. Therefore, the above-mentioned strategy is not applicable to sweetpotato, and a novel method that takes full advantage of genome-wide homologous variation between hexaploid sweetpotato and tetraploid wild relatives is required to more fully examine the origin of sweetpotato.

After comparative studies of *Ib*T-DNA insertions, nuclear and chloroplast genome variations, plus haplotype-based phylogenetic analysis, we revealed the origin of sweetpotato, pointed out the progenitors’ contributions to germplasm in term of T-DNAs, chloroplast genomes and subgenomes. We also identified biased gene conversion events between sweetpotato subgenomes based on homologous haplotypes. Moreover, we provided new insight in the role that selection played in the domestication process of cultivated sweetpotato and identified useful candidate genes for future breeding and genetic engineering efforts, and evolutionary studies. In addition, the identification of the presumptive progenitors will accelerate work towards the generation of artificial hexaploids in the genus *Ipomoea*. Taken together, the results of the present study shed light on the evolution of sweetpotato and pave the way for the genetic improvement of sweetpotato.

## Results

### Phylogeny and population structure of sweetpotato and its wild relatives

To investigate the phylogenetic relationship of sweetpotato and its wild relatives, we analyzed 23 sweetpotato cultivars/landraces and all putative genetic donors of sweetpotato, representing a wide range of taxonomic groups, geographic distribution, and ploidy levels (Fig. 1 and Supplementary Table 1). As diploid relatives, we included five accessions of *I. trifida*, the species that most likely to be the diploid progenitor of sweetpotato, and two wild relatives *I. triloba* and *I. tenuissima*. We also sampled the 45 wild tetraploid accessions which previously reported as the possible progenitor of sweetpotato, including *I. tiliacea, I. aequatoriensis, I. batatas* var. *apiculate, I. tabascana, I. batatas* 4x and possible hybrid *Ipomoea* accessions.

**Fig. 1.**
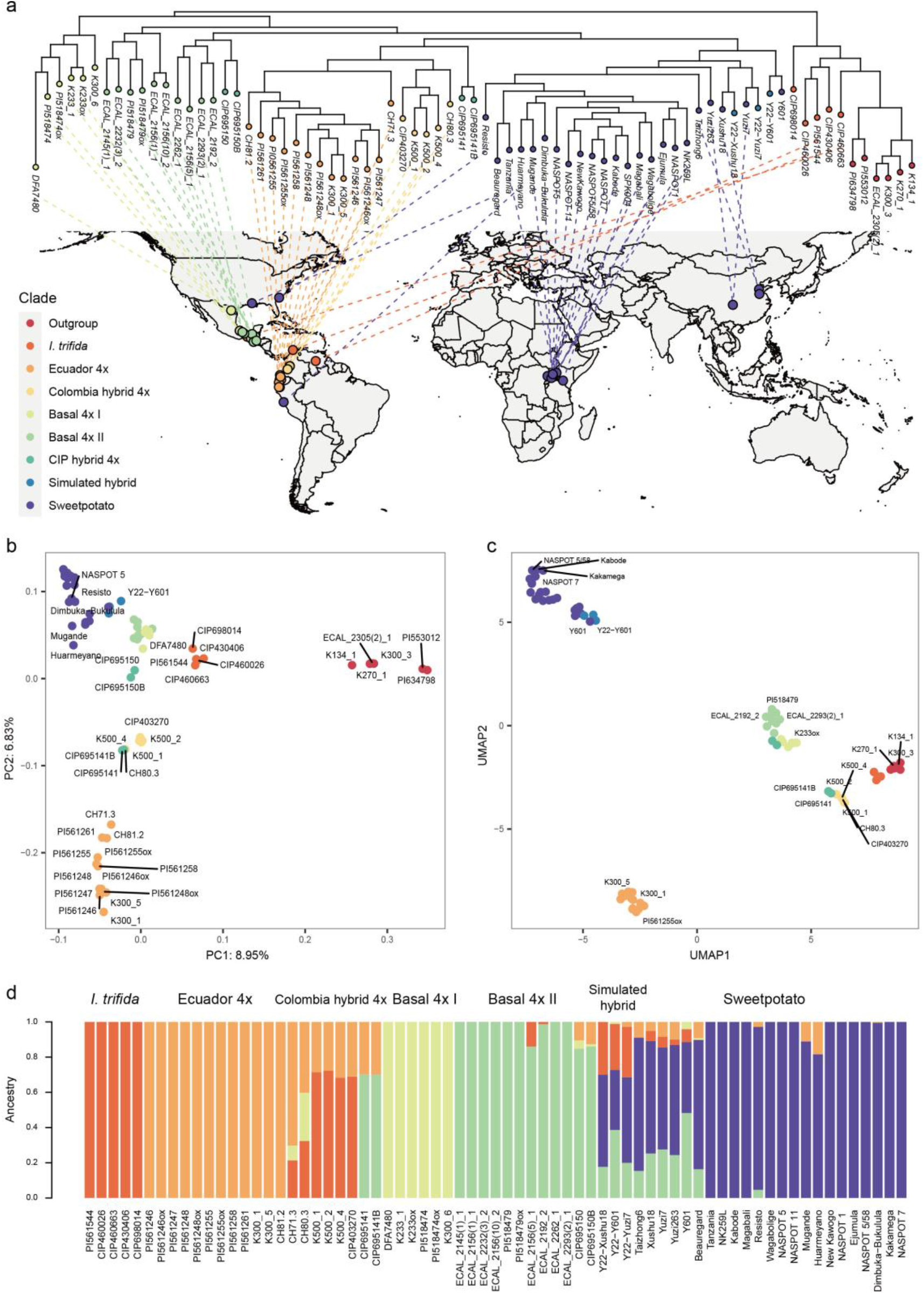
Phylogeny and population structure of sweetpotato and its wild relatives. **a**, Phylogenetic tree based on the genome-wide variants demonstrates the relationships between sweetpotato cultivars/landraces and their wild relatives, were inferred using the maximum likelihood method. All nodes are 100% supported by bootstrap values. The clades are color-coded and colors of **b**-**d** are consistent. The dashed lines link the phylogenetic position on the tree with the geographic location on the map for each accession. The accessions with unknown geographic location are not linked to map. **b**, PCA of sweetpotato and its relatives. The proportions of variance explained by PC1 and PC2 are illustrated on axises. **c**, UMAP using first three principal components. Dot colors are the same as in **a. d**, Population structure analysis of sweetpotato and its close wild relatives for K = 5.

Phylogenetic analyses (Fig.1a and Supplementary Fig. 1) based on 6,326,447 whole genome variations revealed that the diploid *I. trifida* and outgroup species, including diploid *I. triloba, I. tenuissima* and tetraploid *I. tiliacea*, form the basal clade in the phylogeny. The basal 4x lineages, including *I. batatas* var. *apiculate* (basal 4x I clade), *I. batatas* 4x and *I. tabascana* (basal 4x II clade), resides at the base of a large lineage composed of sweetpotato cultivars and a monophyletic tetraploid lineage. The monophyletic tetraploid lineage consists of two monophyletic clades, including tetraploid *I. aequatoriensis* from Ecuador (Ecuador 4x clade) and tetraploid hybrids from Colombia (Colombia hybrid 4x) (Fig.1a). Sweetpotato cultivars/landraces form a sister monophyletic lineage to tetraploid lineage consists of Ecuador 4x clade and Colombia hybrid 4x. Principal component analysis (PCA), uniform manifold approximation and projection (UMAP) and admixture-based analyses clustered all accessions into six main groups, i.e., outgroup, *I. trifida*, Ecuador 4x, Colombia hybrid 4x, basal 4x and sweetpotato (Fig.1 b-d and Supplementary Fig. 2-b,d-e, Fig. 3a). These results are consist with the phylogenetic clades of sweetpotato and its wild relatives. The detailed classification of all samples are provided in Supplementary Table 1.

There are two speculations about the relationship between the basal 4x and sweetpotato. The first one considers the basal 4x as the tetraploid progenitor of sweetpotato ^22^, whilst the second one treats the basal 4x as hybrid offsprings between sweetpotato and *I. trifida* ^23^. In current study, we simulated three tetraploid hybrids of *I. trifida* and sweetpotato by randomly sampling reads from the closest accession of *I. trifida* (CIP698014) related to sweetpotato, and three sweetpotato cultivars at a ratio of 1:3 and integrating the sampled reads respectively. These simulated tetraploid hybrids fell into the sweetpotato clade or cluster in all analyses, and separated from other wild tetraploid relatives (Fig.1 a-d), suggesting that all the wild tetraploid relatives are not hybrids of *I. trifida* and sweetpotato. The Colombia hybrid 4x lies in the middle of *I. trifida*, basal 4x and Ecuador 4x clades (Fig.1 b-c), and the population structure also supports that Colombia hybrid 4x is likely to be the hybrids of *I. trifida*, basal 4x or Ecuador 4x groups (Fig.1 d).

### Horizontal transferred *Ib*T-DNAs reveal two progenitors of sweetpotato

Because the genomes of almost all sweetpotato cultivar/landrace contain horizontally transferred *Ib*T-DNA1 and/or *Ib*T-DNA2 sequences from *Agrobacterium* spp. ^24,25^, *Ib*T-DNAs are most likely inherited from the progenitors of sweetpotato. Therefore, *Ib*T-DNAs serve as natural genetic markers to track the progenitors of sweetpotato ^24^. *I. tenuissima* is the only diploid species contains *Ib*T-DNA1 in this study, but its *Ib*T-DNA1 sequence is very different from those of sweetpotato (Fig.2a and Supplementary Table 1). As for the tetraploid relatives, six accessions of the basal 4x clade (*Ipomoea batatas* 4x and *I. batatas* var. *apiculate*) and three hybrid tetraploid accessions (CIP695141, CIP695150B and CIP403270) contain *Ib*T-DNA1 (Fig.2a and Supplementary Table 1). One of sweetpotato progenitors is very likely to be in the basal 4x clade (*I. batatas* 4x and *I. batatas* var. *apiculate*), since it is the only non-hybrid wild tetraploid relative of sweetpotato containing *Ib*T-DNA1. The phylogeny and structure variations of *Ib*T-DNA1 sequences also indicate sweetpotato resemble the basal 4x clade (Fig.2a). The *Ib*T-DNA1 sequences of several accessions belong to the basal 4x clade are partially covered with sequencing reads (Fig.2a), demonstrating a process that the *Ib*T-DNA1 of the basal 4x clade has been gradually lost in these accession. This explains why the other accessions of the basal 4x clade demonstrate closer relationship with sweetpotato, but does not contain *Ib*T-DNA1 insertion.

**Fig. 2.**
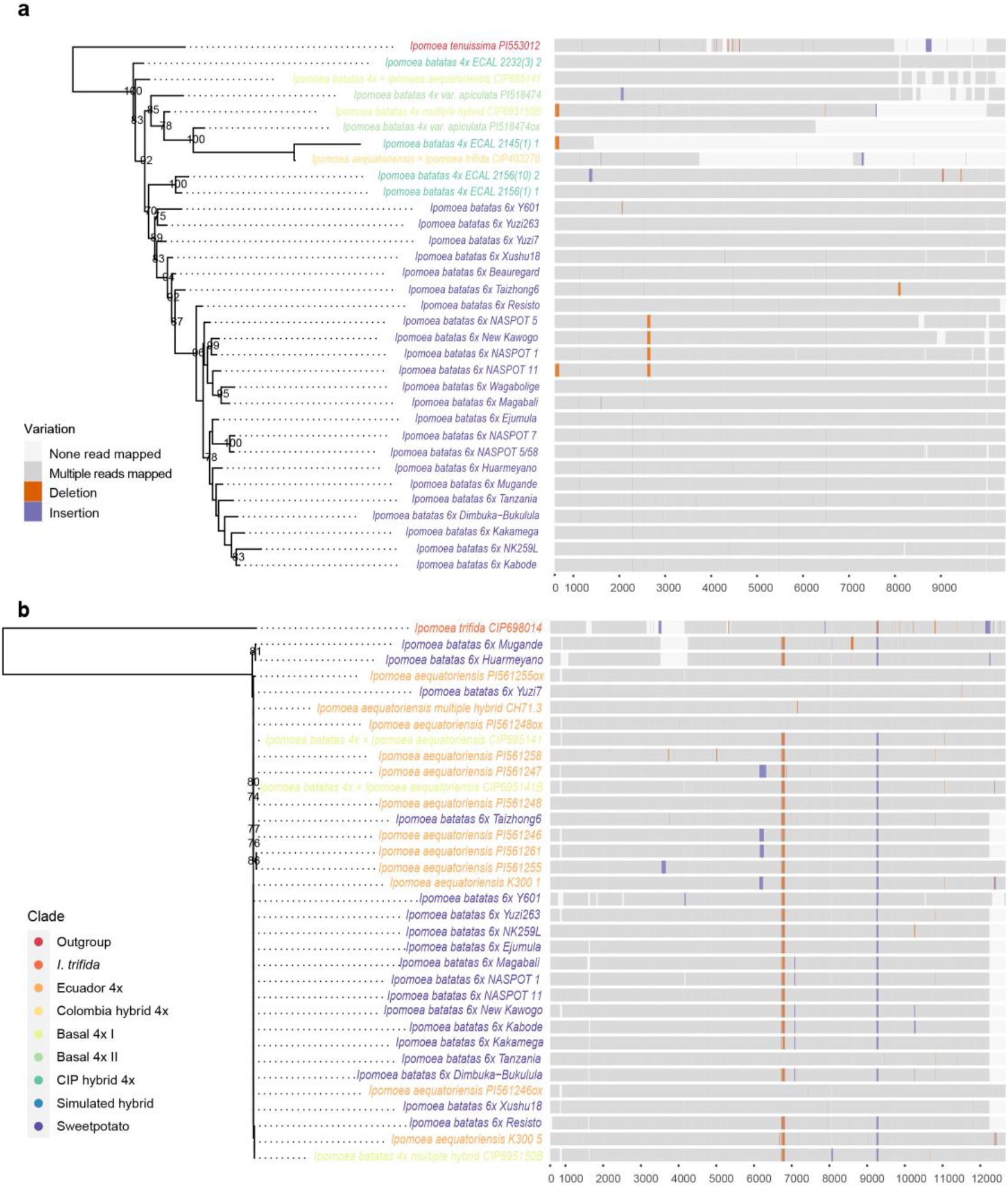
Phylogeny and structure of *Ib*T-DNAs of sweetpotato and its wild relatives. **a**, Maximum likelihood tree of *Ib*T-DNA1 based on variants from positive accessions. Bootstrap values >70% are shown at nodes. The structures of *Ib*T-DNA1 in positive accessions are illustrated on the right. The regions without any reads mapped are likely deletions, which are colored in light gray. The indels are also color-coded. **b**,Maximum likelihood tree of *Ib*T-DNA2 based on sequence variants from positive accessions. Bootstrap values >70% are shown at nodes. The structures of *Ib*T-DNA2 in positive accessions are illustrated on the right.

*I. trifida* was previously considered as the diploid progenitor of sweetpotato. We identified six *Ib*T-DNA2 positive accessions after screening 37 accessions of *I. trifida* (Fig.2b and Supplementary Table 1). However, *Ib*T-DNA2 sequences of all six positive accessions form a sister lineage of sweetpotato, as revealed by phylogeny and *Ib*T-DNA2 structure (Fig.2b and Supplementary Figure 4). Meanwhile, accessions of tetraploid *I. aequatoriensis* (belong to the Ecuador 4x clade) and three artificially hybrid tetraploid accessions (CIP695141, CIP695141B and CIP403270) also contain *Ib*T-DNA2 insertion (Fig. 2b and Supplementary Table 1). Furthermore, *Ib*T-DNA2 sequences of accessions of *I. aequatoriensis* resemble those of sweetpotato since they fall in the same lineage with sweetpotato (Fig.2b). Therefore, *I. aequatoriensis* is more likely related to the progenitor which passed the *Ib*T-DNA2 to sweetpotato.

### The subgenome origins of sweetpotato revealed by haplotype-based phylogenetic analysis (HPA)

To figure out which subgenome contributed by each progenitor, the relationships between sweetpotato and each possible progenitors are informative. Considering the dosage effect, the progenitor contributed four copy of B_2_ subgenome (tetraploid progenitor) is closer to sweetpotato than the progenitor contribute two copy of B_1_ subgenome (diploid progenitor) (Fig. 3a). The basal 4x clade is more closely related to sweetpotato than *I. aequatoriensis* in PCA (only PC1 vs PC2), UMAP plots (Fig.1b-c) and genome-wide nucleotide diversity (Supplementary Fig. 5-6), although *I. aequatoriensis* is the sister group of sweetpotato (Fig.1a). However, these analyses treated the hexaploid sweetpotato and tetraploid relatives as diploid, which artificially decreased the allelic variations of polyploids.

**Fig. 3.**
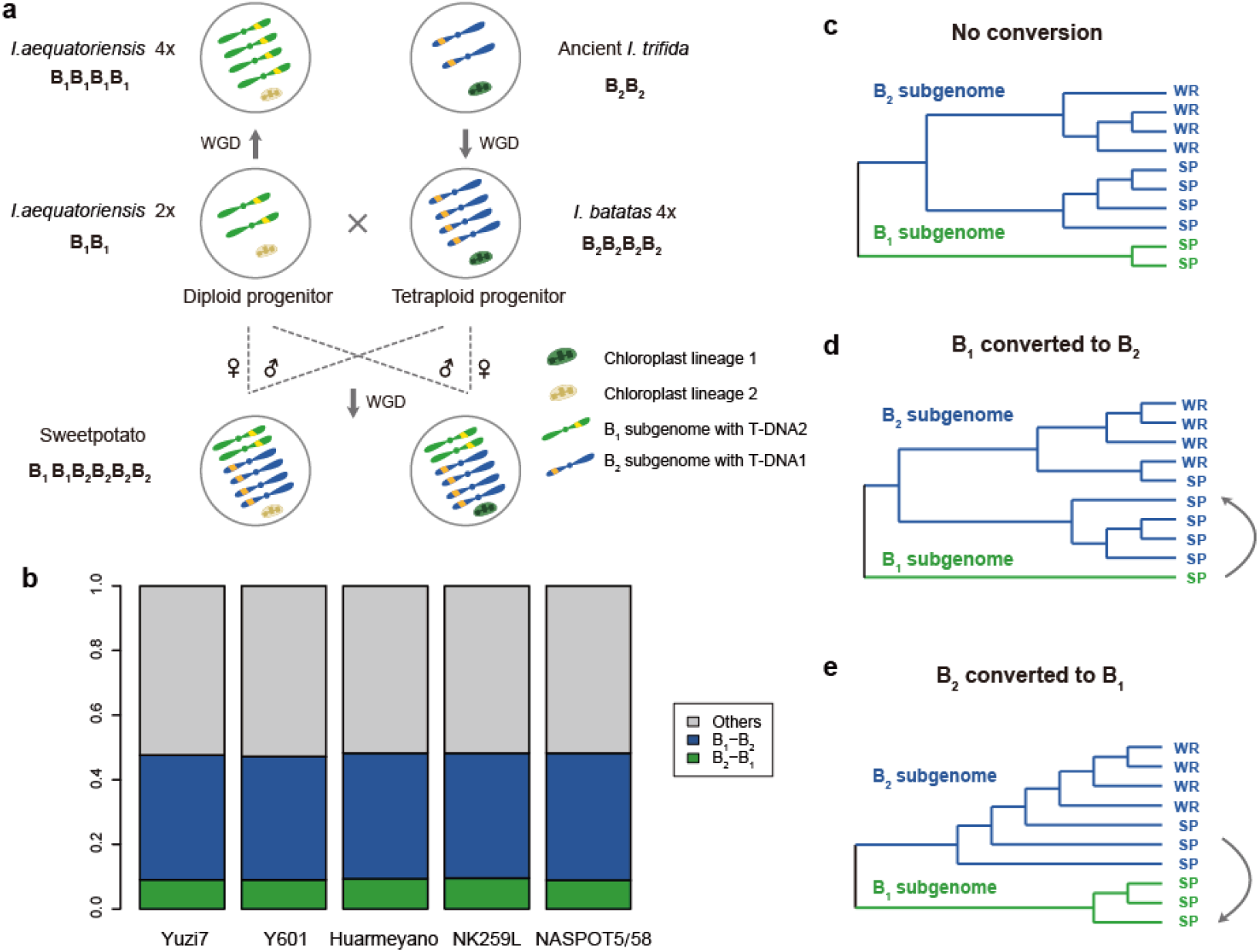
Origin hypothesis of sweetpotato and gene conversions between sweetpotato subgenomes. **a**, origin hypothesis of sweetpotato. The diploid *I. aequatoriensis* is likely to be the diploid progenitor, contributed the B_1_ subgenome, *Ib*T-DNA2 and lineage 2 type of chloroplast genome to sweetpotato. The tetraploid progenitor of sweetpotato is *I. batatas* 4x (derived from duplication of ancient *I. trifida*), donating the B_2_ subgenome, *Ib*T-DNA1 and lineage 1 type of chloroplast genome. Sweetpotato derived from the reciprocal cross between the diploid and tetraploid progenitors and a subsequent whole genome duplication. **b**, Gene conversion ratios in five hexaploid sweetpotato cultivars/landraces using the closest natural accession (ECAL_2262_1) resembling the tetraploid progenitor as reference. B_1_ – B_2_, gene conversion events from the B_1_ to the B_2_ subgenome. B_2_ - B_1_, conversion events from B_2_ to B_1_ subgenome. Others, other scenarios, including no conversion and scenarios that could not be resolved. **c-e**, Examples of tree topologies under the scenarios of no conversion (**c**), B_1_ to B_2_ gene conversion (**d**), and B_2_ to B_1_ gene conversion (**e**). The B_1_ subgenome is shown in green and the B_2_ subgenome in blue. SP, sweetpotato. WR, wild relative (tetraploid progenitor).

To reveal the accurate relationship between sweetpotato and its progenitors, we developed a HPA pipeline which uses homologous haplotypes of polyploid to conduct high-throughput phylogenetic analyses (Supplementary Fig. 7). First, we independently phased the genome sets of three representative hexaploid sweetpotato cultivar and 38 tetraploid accession. As for the representative sweetpotato cultivars, we chose representative cultivar from phylogenetic lineages in the sweetpotato phylogeny (Fig.1; Supplementary Fig. 1), i.e., Huameyano, NK259L, and Yuzi7. Each cultivar was used to extract the syntenic haplotype block with each tetraploid accession. We obtained 439,555-760,769 haplotype blocks in the three sweetpotato cultivars (Supplementary Table 2; Supplementary Fig. 8a) and 380,895-1,007,206 haplotype blocks in the 38 tetraploid accessions (Supplementary Table 3; Supplementary Fig. 8b). Second, we extracted the syntenic haplotype blocks shared between each sweetpotato cultivar and each tetraploid accession by comparing their genomic positions. In doing so, we identified 606,246-1,154,274 syntenic haplotype blocks (Supplementary Table 4; Supplementary Fig. 9). Third, we removed (i) redundant syntenic haplotype blocks that had overlapping regions with other blocks, and (ii) those blocks that consist of very short sequences (less than 20 bp). Ultimately, 412,632-866,522 syntenic haplotype blocks were extracted, which accounted for 28.2-41.7% of the sweetpotato genome (Supplementary Table 5; Supplementary Fig. 10).

The previously identified syntenic haplotype blocks between each sweetpotato cultivar and each tetraploid accession were used to perform phylogenetic reconstructions independently. The phylogenetic trees were inferred by two methods: Unweighted Pair-Group Method with Arithmetic Mean (UPGMA) and maximum likelihood (ML). We calculated the monophyletic ratio and the Nsp-Nwr distance to measure the relationship between the investigated tetraploid accession and the representative hexaploid sweetpotato (Supplementary Fig. 7d). The monophyletic ratio is defined as the proportion of trees in which sweetpotato haplotypes forming a monophyletic clade (Supplementary Fig. 7d). The Nsp-Nwr distance is defined as the tree branch length between the most recent common ancestor (MCRA) node of sweetpotato haplotypes (i.e., Nsp) and the MCRA node of the tetraploid accession (i.e., Nwr) (Supplementary Fig. 7d). PI index is a coefficient that calculated the difference between haplotype nucleotide diversity of sweetpotato and the tetraploid accession (Supplementary Fig. 7d). For mentioned three indices, smaller value indicates a closer relationship between the investigated tetraploid accession and the hexaploid sweetpotato. To increase accuracy, we only included trees that had the same monophyletic judgement by both tree-building methods, and these trees were used to calculate the monophyletic ratio and Nsp-Nwr distance. Among all syntenic haplotype blocks, the 6:4 data set (composed of six haplotypes of sweetpotato and four haplotypes of tetraploid accessions) produced the most robust results, since results of the 6:4 data set are consist based on the three indices using the three sweetpotato cultivars (Fig. 4a-c and Supplementary Fig. 11-25).

**Fig. 4.**
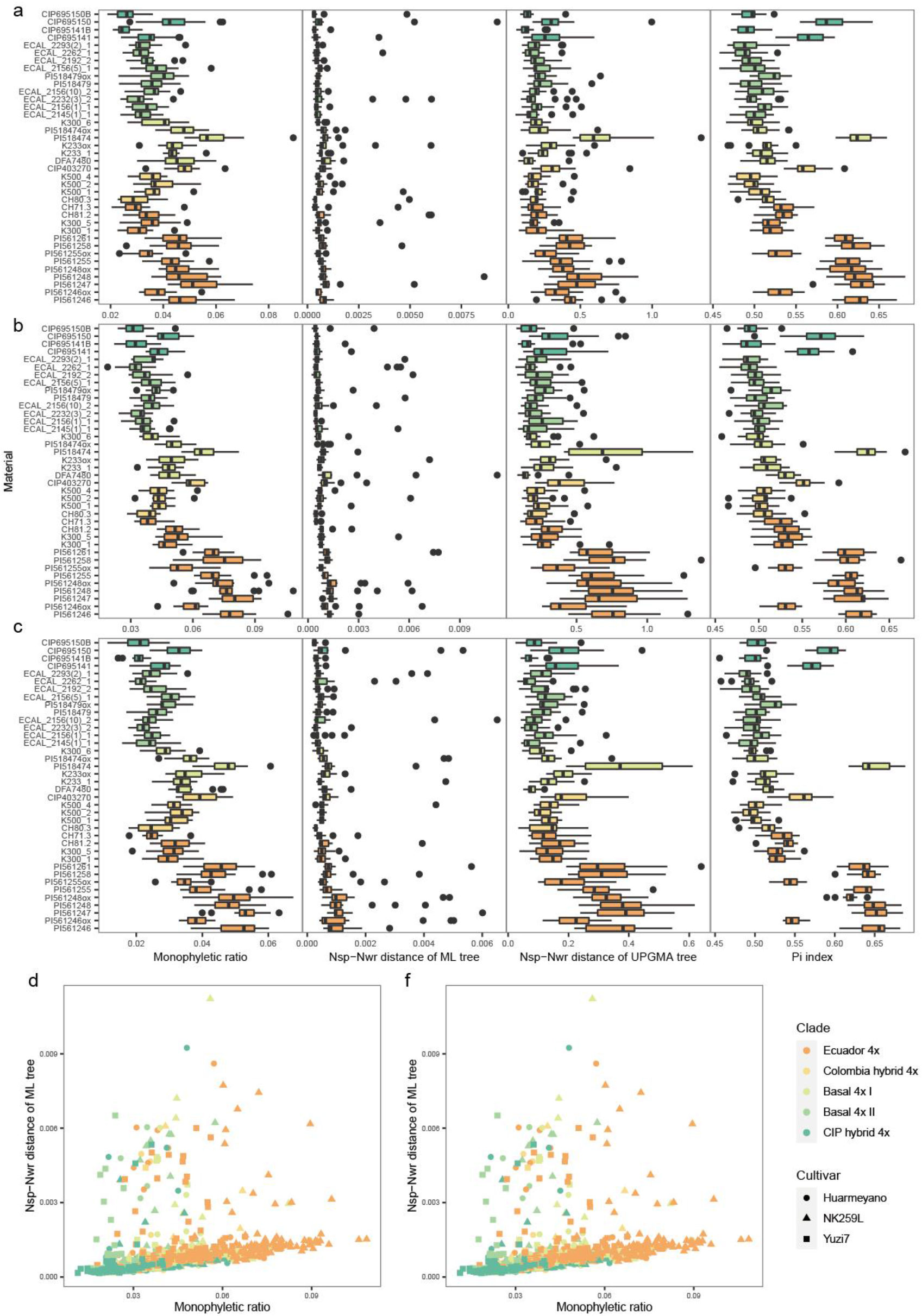
Relationships between sweetpotato cultivars and tetraploid accessions as revealed by HPA. Boxplots of the monophyletic ratios, the Nsp-Nwr distances of two methods, and PI index of 15 chromosomes among 38 tetraploid accessions. **a**, the results of cultivar Huarmeyano. **b**, the results of cultivar NK259L. **c**, the results of cultivar Yuzi7. Monophyletic ratio, the proportion of trees in which sweetpotato haplotypes forming a monophyletic clade. Nsp-Nwr distances, the tree branch length between the most recent common ancestor (MCRA) node of sweetpotato haplotypes (i.e., Nsp) and the MCRA node of the tetraploid accession (i.e., Nwr). PI index, a coefficient that calculated the difference between haplotype nucleotide diversity of sweetpotato and the tetraploid accession.

HPA provides a better resolution and relatively consistent results to resolve the relationship between sweetpotato and tetraploid relatives. All three indices of HPA show that, among the non-hybrid tetraploid relatives, the basal 4x clade is the closest relatives of sweetpotato and *I. aequatoriensis* is the farthest tetraploid relative (Fig. 4a-f). Besides, HPA also enables to resolve that *I. batatas* 4x and *I. tabascana* (the basal 4x II clade) is closer related to sweetpotato than *I. batatas* var. *apiculate* (the basal 4x I clade) (Fig. 3). Therefore, *I. batatas* 4x is most likely to be the tetraploid progenitor, which contributed B_2_ subgenome to sweetpotato (Fig. 3a). The accession ECAL_2262_1 has shown the closest relationship with sweetpotato, although it does not contain *Ib*T-DNA1 insertion. ECAL_2262_1 might have gradually lost *Ib*T-DNA1 insertion completely as the process demonstrated in other accessions of *I. batatas* 4x (Fig. 2a). It has not escaped our notice that *I. aequatoriensis*, another potential progenitor species revealed by *Ib*T-DNA2, is most likely related to the diploid progenitor, which contributed B_1_ subgenome to sweetpotato (Fig. 3a). Therefore, the diploid *I. aequatoriensis* is very likely to be the diploid progenitor of sweetpotato.

CIP hybrid 4x group is most closely related to sweetpotato (Fig. 4a-f). CIP hybrid 4x are artificial hybrids between tetraploid relatives. *I. batatas* 4x and *I. aequatoriensis* are involved in the pedigrees of CIP hybrid 4x accessions. The closest relationship with sweetpotato is probably because CIP hybrid 4x shares the similar genetic background with sweetpotato, and supports *I. batatas* 4x, *I. aequatoriensis* are the progenitors of sweetpotato or related to sweetpotato progenitors. The genetic backgrounds of CIP hybrid 4x are described in the Supplementary Note.

### Chloroplast genome confirmed the identification of two sweetpotato progenitors

The chloroplast haplotypes of sweetpotato are divided into two lineages, i.e., lineage 1 and lineage 2, and the haplotypes of two progenitors are resided in the two lineages (Fig. 5 and Supplementary Fig. 27). The chloroplast haplotypes of *I. batatas* 4x is nested in the lineage 1 of sweetpotato, and the closest individuals are five accessions of *I. batatas* 4x (ECAL_2156(1)_1, ECAL_2156(10)_2, ECAL_2192_2, ECAL_2262_1 and ECAL_2293(2)_1) (Fig. 5 and Supplementary Fig. 27). Other accessions of *I. batatas* 4x, *I. tabascana, I. batatas* var. *apiculate* and Colombia hybrid 4x also fall into lineage 1 but are relatively far related to sweetpotato haplotypes (Fig. 5 and Supplementary Fig. 27). Therefore, chloroplast haplotypes also support *I. batatas* 4x is the progenitor of sweetpotato. Besides, as the species resembles the diploid progenitor of sweetpotato, *I. aequatoriensis* is the only non-hybrid species nested in the lineage 2 of sweetpotato haplotypes (Fig. 5 and Supplementary Fig. 27). Therefore, the two chloroplast haplotype lineages of sweetpotato are likely inherited from its two progenitors directly (Fig. 3a). The haplotypes of *I. trifida* are relatively far from sweetpotato than the two progenitors (Fig. 5 and Supplementary Fig. 27), which indicates the extant *I. trifida* may not be the diploid progenitor of sweetpotato.

**Fig. 5.**
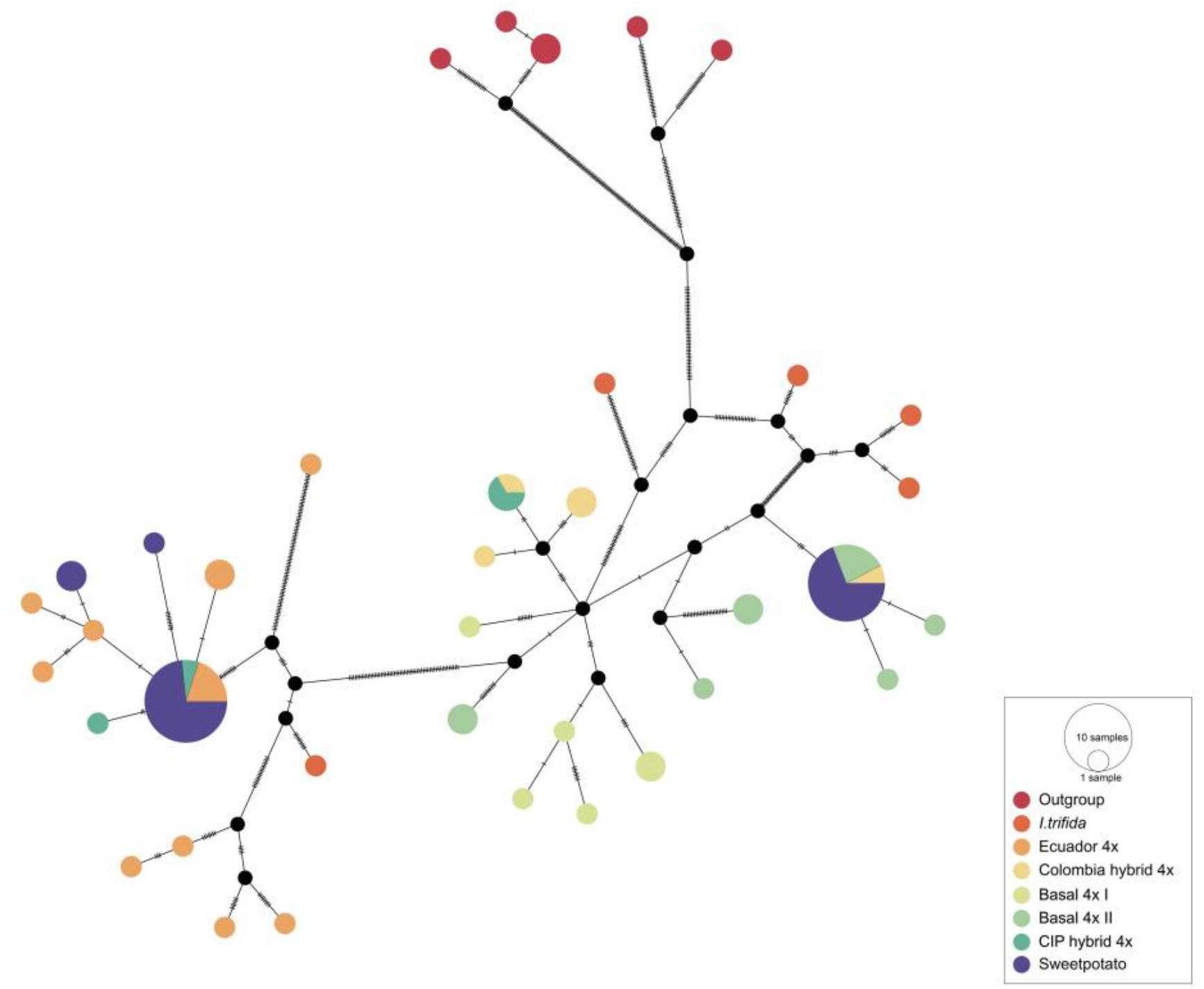
The phylogenetic network of chloroplast genomes of sweetpotato and its wild relatives. The network inferred using TSC network based on chloroplast genome. Circle size is proportional to the frequency of a haplotype across all populations. Each line between two haplotypes represents a mutational step. Number of short lines at the middle of the edges indicates the number of hypothetical missing haplotypes. The solid black dot means existing of unsampled haplotypes or extinct ancestral haplotypes. The clades are color-coded.

### Gene conversion between sweetpotato subgenomes

Gene conversion in polyploids refers to sequence exchanges between homologous genes from different subgenomes, in which one progenitor allele overwrites another ^36–38^. The sweetpotato genome is comprised of two B_1_ and four B_2_ subgenomes (B_1_B_1_B_2_B_2_B_2_B_2_). Subgenomes B_1_B_1_ were donated by the diploid progenitor and subgenomes B_2_B_2_B_2_B_2_ by the tetraploid progenitor ^20^ (Fig. 3a). If no conversion events occurred, each syntenic haplotype block should have two copies of the B_1_ subgenome from sweetpotato, four copies of the B_2_ subgenome from sweetpotato, and four copies of the B_2_ subgenome from *I. batatas* 4x (Fig. 3a, c). If a gene were converted between B_1_ and B_2_ subgenomes, the copy numbers of subgenomes and tree topology should deviate from the standard 2:8 ratio between B_1_ and B_2_ in the hexaploid sweetpotato and *I. batatas* 4x (Fig. 3c-e). To detect possible gene conversion events, we first filtered those syntenic haplotype blocks and use blocks in gene regions with six haplotypes of sweetpotato and four haplotypes of *I. batatas* 4x. Finally, 13,535-27,867 homogeneous haplotype blocks in gene regions of sweetpotato cultivars and the closest *I. batatas* 4x accession (ECAL_2262_1), which resembles the tetraploid progenitor, are obtained to identify gene conversion events between subgenomes (Supplementary Table 6). The analysis pipeline has been illustrated in Supplementary Fig. 28 and described in detail in Supplementary Note. Using five sweetpotato cultivars as references, 47.1-48.3% of gene regions in sweetpotato showed evidence of conversion between subgenomes (Fig. 3b; Supplementary Table 6). We found that B_1_ to B_2_ subgenome gene conversions (38.1-39.3%) were much more common than B_2_ to B_1_ conversions (8.9-9.6%) (Fig. 3b; Supplementary Table 6). This was to be expected, as gene conversion is known to be a copy number-dependent process ^39^.

## Discussion

Understanding the genetic origin of crops is vital for breeding and genetic engineering efforts, and is particularly important to all genetic improvement strategies involving wild relatives. The origin of sweetpotato is still the subject of fierce debate. Competing hypotheses have been put forward proposing that sweetpotato is an autopolyploid, a segmental allopolyploid, or an allopolyploid ^8,9,16,18,19,22,23,40,41^. The genetic origin of sweetpotato has remained unresolved because of the high complexity of the genome, due to its hexaploid nature and high degree of heterozygosity ^5,26^. In addition, the two progenitors of sweetpotato are genetically closely related, thus adding to the difficulties in distinguishing the subgenomes of sweetpotato. The half-phased genome sequence of sweetpotato has identified that two sets of chromosomes contributed by a diploid progenitor and other four sets of chromosome came from a tetraploid progenitor ^5^, and confirmed the B_1_B_1_B_2_B_2_B_2_B_2_ genome architecture that has been revealed by earlier cytogenetic studies ^19,20^. Therefore, both genomic and cytogenetic analyses suggest that sweetpotato arose from a cross between a diploid progenitor and a tetraploid progenitor.

Here, we propose an origin hypothesis of sweetpotato (Fig. 3a), which meets all known genetic features of sweetpotato, including *Ib*T-DNA insertions, nuclear variations and chloroplast genome. The diploid *I. aequatoriensis* is likely to be the diploid progenitor, contributed the B_1_ subgenome, *Ib*T-DNA2 and lineage 2 type of chloroplast genome to sweetpotato. The tetraploid progenitor of sweetpotato is *I. batatas* 4x (probably derived from duplication of ancient *I. trifida*), donating the B_2_ subgenome, *Ib*T-DNA1 and lineage 1 type of chloroplast genome. Sweetpotato derived from the reciprocal cross between the diploid and tetraploid progenitors and a subsequent whole genome duplication. This hypothesis provides a reasonable explanation about the origin of two subgenomes, *Ib*T-DNA insertions and two lineages of chloroplast genome within the cultivated sweetpotato.

For a long time, *I. trifida* is considered as the diploid progenitor of sweetpotato, because it is the closest diploid species of sweetpotato revealed by DNA sequence and cytogenetic evidences ^7,8,14,15^. However, the *Ib*T-DNA2 sequences of *I. trifida* are very different from those of sweetpotato. Furthermore, the extant accessions of *I. trifida* are relatively far from sweetpotato compared with the two progenitors, as elucidated in this and previous studies ^22,23^. All the facts suggest that the extant *I. trifida* is unlikely to be the direct diploid progenitor of sweetpotato, or the *I. trifida* individuals resemble the diploid progenitor have not be found yet. *I. aequatoriensis* is a recently named autotetroploid species, which was identified as the tetraploid progenitor of sweetpotato in previous study ^23^. The *Ib*T-DNA2 sequences of *I. aequatoriensis* are very similar with insertions within sweetpotato genomes. Besides, both the nuclear and chloroplast variations indicate *I. aequatoriensis* is closer related to sweetpotato than the extant *I. trifida*, as revealed in this and previous studies ^22,23^. Therefore, diploid *I. aequatoriensis* is very likely related to the diploid progenitor of sweetpotato, and that’s why two species form a close sister relationship in the nuclear phylogeny.

Except for the hybrids, *I. batatas* 4x is the closest tetraploid species related to sweetpotato, as revealed in this study and previous reports ^22,23^. Meanwhile, *Ib*T-DNA1 sequences within *I. batatas* 4x resemble *Ib*T-DNA1 sequences in sweetpotato very closely. Furthermore, chloroplast genomes of *I. batatas* 4x fall into the lineage 1 of sweetpotato. These facts support *I. batatas* 4x to be the tetraploid progenitor of sweetpotato. However, *I. batatas* 4x was previously identified as the hybrid between *I. trifida* and sweetpotato ^23^. The key to confirm the tetraploid progenitor is to establish an effective standard to distinguish the possible tetraploid progenitor and the hybrid offspring ^21^. Therefore, we simulated three hybrids between sweetpotato and *I. trifida* and they clustered within the sweetpotato clade instead of the basal 4x clade (*I. batatas* 4x). The population structure analysis also supports *I. batatas* 4x is not hybrid between sweetpotato and *I. trifida* (Fig. 1d). The close relationship between sweetpotato and *I. batatas* 4x is attributable to the fact that *I. batatas* 4x is the tetraploid progenitor and contribute two thirds of chromosomes to sweetpotato. The species *I. tabascana* with only a single collection ^42^ was also suggested to be hybrid between sweetpotato and *I. trifida* ^23,41^. However, it falls in the basal 4x clade based on the nuclear phylogeny and nested in the lineage 1 of chloroplast network, closed to *I. batatas* 4x chloroplast haplotypes. Therefore, *I. tabascana* belongs to *I. batatas* 4x and unlikely to be hybrid offspring between sweetpotato and *I. trifida. I. tiliacea* was another species identified as possible tetraploid progenitor ^1^. Hence, we also included this species into analyses. However, both the nuclear and chloroplast variations indicate *I. tiliacea* is far related to sweetpotato. Meanwhile, *Ib*T-DNAs are not found in the four accessions of *I. tiliacea*. Therefore, *I. tiliacea* is unlikely to be the tetraploid progenitor of sweetpotato.

None of previous origin hypotheses could explain the formation of the two distinct lineages of sweetpotato in the chloroplast phylogenies. The first hypothesis suggested the asymmetrical hybridization between diploid *I. trifida* and original hexaploid sweetpotato result in chloroplast capture from *I. trifida* ^8,23^. However, this explanation ignored the asymmetrical hybridization between a diploid and hexaploid will decrease the ploidy level of offsprings from hexaploid to tetraploid. The second hypothesis is the hybridization between sweetpotato and *I. trifida* produced a new allotetraploid entity, and subsequently hybridized with *I. trifida* and formed a new hexaploid form. Then, the newly formed hexaploid repeatedly crossed with the original hexaploid *I. batatas*, progressively losing the *I. trifida* component of its nuclear genome while maintaining a *trifida*-like chloroplast ^8^. This explanation is tedious since two hybridization events to form a new allotetraploid and repeatedly asymmetrical hybridization are both required. Furthermore, chloroplast genomes of *I. trifida* form an independent lineage distinct from the two lineages of sweetpotato. Therefore, considering chloroplast genome, the extant *I. trifida* could not be the progenitor of sweetpotato. However, our work provides a simpler explanation that the two types of sweetpotato chloroplast genomes are inherited from its two progenitors directly, which is supported by chloroplast phylogeny. During the formation of sweetpotato, the two progenitors crossed reciprocally and passed the two type of chloroplast genomes to sweetpotato.

Accurate relationship between sweetpotato and tetraploid relatives is the key to identify the subgenome origin of sweetpotato. Using the routine phylogenetic and population structure methods, all polyploids have to be treated as diploids to meet the data format required by current available softwares. This procedure artificially decreased the nucleotide diversity of polyploids, and unavoidably resulted in uncertainty in the conclusions that could be drawn. This represents a common problem in studies on the origin of polyploid species that rely on consensus variation. To solve this problem, we developed a HPA pipeline that takes full advantage of homologous variation while maintaining the true nucleotide diversity of the polyploid species. This new pipeline authenticated the result of the consensus genome-wide variation analysis. Furthermore, HPA provides a better resolution, results in more accurate relationship between sweetpotato and various tetraploid relatives. We successfully identified the two progenitors of sweetpotato and the closest accessions related to the two progenitors. The closest accessions related to the diploid progenitor is the tetraploid accession of *I. aequatoriensis* (PI561255) from Ecuador, and the closest accessions related to the tetraploid progenitor is a Mexican accession of *I. batatas* 4x (ECAL_2262_1). Since the two probable progenitors are distributed in Central America, including South Mexico, Guatemala, Ecuador and Venezuela, Central America is likely to be the original place where sweetpotato formed naturally. Unfortunately, the unambiguous diploid form of *I. aequatoriensis* (the diploid progenitor) has not been discovered yet. Considering *Ib*T-DNA2 sequence has been proved to be an effective marker to identify the diploid progenitor of sweetpotato, it is necessary to conduct a wider survey and full examination of diploid species in Central America to search sweetpotato-type of *Ib*T-DNA2 sequence, to ultimately discover the diploid progenitor of sweetpotato. Quispe-Huamanquispe (2019) ^24^ provides a practical methods and demonstrated screening only one gene (*ORF13*) and simple phylogenetic analysis will be effective enough to preliminarily identify the diploid progenitor of sweetpotato. Sweetpotato breeders have been working for decades towards generating artificial hexaploids from diploid and tetraploid wild relatives of sweetpotato ^58^. The discovery of the two progenitors of sweetpotato and their extant closest accessions not only contributes to our understanding of the genetics of sweetpotato, but also provides a critical natural resource for future breeding programs.

Another important application of HPA lies in the use of homologous haplotypes to detect gene conversion between subgenomes. In sweetpotato, almost half of the gene regions show evidence of conversion between subgenomes. Taking advantages of phased haplotypes, the identified gene conversion events between subgenomes also shed light on the evolution and domestication of hexaploid sweetpotato. B_1_ to B_2_ conversion events are approximately 3-times more frequent than B_2_ to B_1_ conversions (Fig. 3b). Rampant gene conversion and conversion biases increase genome complexity in sweetpotato and may suggest an important role for gene conversion in genome evolution and domestication of sweetpotato. Subgenome-biased conversion has been reported in several allopolyploid crop plants including cotton, canola, peanut and strawberry ^35,37,43,44^. However, the molecular mechanisms underlying the conversion bias are largely unknown. In the case of sweetpotato, the dosage effect (of the tetraploid B_2_ versus the diploid B_1_ genome) may explain the more prevalent conversion of B_1_ alleles to B_2_ alleles. Because gene conversion is known to be a copy number-dependent process ^39^. But the phased haplotypes are still fragmental and short (Supplementary Table 5 and Supplementary Fig. 26), long haplotype phasing using nanopore or PacBio sequences or fully-phased genome assembly are very necessary to accurately confirm the subgenome origin and conversion in the future. The new knowledge on sweetpotato genomics and domestication revealed in this study will contribute to this goal and aid future breeding and genetic engineering approaches in this important staple crop.

## Methods

### Plant materials

Seven diploid wild relatives of sweetpotato (including five accessions of *I. trifida*, one accession of *I. triloba* and one accession of *I*. sp), 42 tetraploid wild relatives of sweetpotato (four accessions of *I. tiliacea*, six accessions of *I. batatas* var. *apiculate*, eight accessions of *I. batatas* 4x, two accessions of *I. tabascana*, twelve accessions of tetraploid *I. aequatoriensis* and ten accessions of hybrids) and 23 sweetpotato cultivars/landraces were utilized in phylogenetic analysis of nuclear and chloroplast genome. Among sweetpotato cultivars/landraces, sequencing data from Taizhong6, Xushu18, Y601, Yuzi263 and Yuzi7 were newly generated in this study. All other data was downloaded from NCBI, including cultivars Tanzania, Beauregard and 16 cultivars in the Mwanga diversity panel (MDP) ^26^. Three tetraploid hybrids of *I. trifida* and sweetpotato were simulated by randomly sampling reads from the accession CIP698014 of *I. trifida*, and three sweetpotato cultivars (Xushu18, Y601, and Yuzi7) at a ratio of 1:3 using seqtk (version 1.3) ^45^. Sampled reads were integrated between *I. trifida* and each sweetpotato cultivar, respectively. We also included 27 accessions of *I. trifida* with low-depth sequenced only for *Ib*T-DNA analyses. Detailed information on the plant materials is given in Supplementary Table 1 and Supplementary Fig. 1-4.

### Resequencing and population analysis

*Variant calling*. The WGS paired-end reads were aligned to the reference sweetpotato genome (https://sweetpotao.com/download_genome.html) using bwa-mem (version 0.7.17) ^46^ and sorted by samtools (version 1.10) ^47^ with the default parameters. Picard (version 2.23.4) ^48^ was used to label PCR duplicates based on the mapping coordinates. 120,369,840 genetic variants including SNPs and INDELs were detected as diploid using the Genome Analysis Toolkit (GATK, version 4.1.8.1) ^49^. About 88% of raw variants were filtered out using VCFtools (version 0.1.17) ^50^ with the following parameters: --minDP 3 --minQ 30 --max-missing 0.8 --maf 0.05. Then, SNPs were filtered based on linkage disequilibrium using PLINK (--indep-pairwise 200 10 0.5) and VCFtools. Finally, a total of 6,326,447 variants were selected and used in in phylogenetic analysis, population genetic diversity analysis and selective sweep detection.

#### Phylogenetic analysis

Vcf2phylip (version 2.7) ^51^ was used to generate a fasta file by concatenating all SNPs from the VCF file and the heterozygous SNPs were degenerated. A phylogenetic tree of sweetpotato cultivars/landraces and wild relatives was reconstructed using IQ-TREE (version 1.6.12)^52^ with 1,000 ultrafast bootstrap replicates. The nucleotide substitution model (GTR+F+I+G4) was selected by IQ-TREE. The phylogenetic tree was rooted with the diploid wild relatives as outgroup and all accessions were plotted onto world map using the R package phytools (version 0.7-70) ^53^.

#### Population structure analysis

PCA was performed using the PLINK (v1.90b6.24) ^54^. The PCA plots were visualized using R package ggplot2 ^55^. The UMAP was performed using R package umap ^56^. The input Plink binary files are transformed from VCFs file using PLINK. Ancestral population stratification was inferred using Admixture (version 1.3.0) ^57^ software, using ancestral population sizes K=1–10.

#### Population genetic diversity

LD decay was calculated using PopLDdecay (v3.27) ^58^ with default parameters. Nucleotide diversity (θπ) were determined for two tetraploid wild relative population (16 accessions of the basal 4x clade and 12 accessions of *I. aequatoriensis*) and the sweetpotato population (23 cultivars/landraces) with VCFtools (version 0.1.17) ^50^ using a 100 kb sliding window with a 10 kb step size.

### Haplotype-based phylogenetic analysis

We developed the HPA pipeline to investigate the relationship of each tetraploid accession to cultivated sweetpotato (Supplementary Fig. 6).

#### Haplotyping

The WGS paired-end reads from the sweetpotato cultivars and *I. batatas* 4x were mapped to the sweetpotato reference genome using bwa-mem (version 0.7.17-r1188). Freebayes (version v1.3.1-17-gaa2ace8) ^59^ was used to call variants (setting −p 6 for sweetpotato and −p 4 for *I. batatas* 4x). Ranbow (version 2.0) ^60^ was used for genome haplotyping.

#### Phylogenetic analysis

The syntenic haplotype blocks between each sweetpotato cultivar and each tetraploid accession were extracted and filtered using HPA pipeline. Sequences within each syntenic haplotype block were aligned by MAFFT (v7.471) ^61^. The UPGMA tree and ML tree for each syntenic haplotype block were reconstructed independently using MEGA-CC (version 10.1.8) ^62,63^ and IQ-TREE respectively. The monophyletic ratio and Nsp-Nwr distance were calculated using HPA pipeline. To increase the accuracy, only those trees which had the same monophyletic judgement by two tree-building methods (trees generated based on the same syntenic block by two methods are both monophyletic or both not monophyletic) were used to calculate monophyletic ratio and Nsp-Nwr distance. The detailed identification procedures are described in the Supplementary Note.

### Gene conversion

The syntenic haplotype blocks that had six haplotypes of sweetpotato and four haplotypes of *I. batatas* 4x, within gene regions, were extracted to detect gene conversion between subgenomes. When ignoring the reverse gene conversion, if there is no gene conversion in a specific syntenic haplotype block, the block is expected to have two B_1_ subgenome haplotypes and four B_2_ subgenome haplotypes from sweetpotato, and four B_2_ subgenome haplotypes from *I. batatas* 4x. The phylogenetic tree should form two clades corresponding to haplotypes of each subgenome. If a gene was converted between subgenomes, the number of haplotypes and the tree topology is expected to vary. Gene conversions were identified based on tree topology (Supplementary Fig. 25). The detailed procedures for determining gene conversion are provided in the Supplementary Note.

### *Ib*T-DNA analysis

#### IbT-DNA detection

The PCR detection of *Ib*T-DNA1 and *Ib*T-DNA2 genes was performed as previously described in Quispe-Huamanquispe, et al. ^24^. The WGS paired-end reads were aligned to *Ib*T-DNA1 and *Ib*T-DNA2 reference sequence (GenBank: KM052616 and KM052617) using bwa-mem (version 0.7.17) ^46^. And the bam files were visualized in IGV (version 2.8.2) ^64^ to check the presence/absence of T-DNA insertions.

#### Phylogenetic and structure variation analysis

Picard (version 2.23.4) ^48^ was used to label PCR duplicates based on the mapping coordinates of the bam files. Genetic variants including SNPs and INDELs were detected as diploid using the Genome Analysis Toolkit (GATK, version 4.1.8.1) ^49^. The methods of phylogenetic analysis is the same as phylogenetic analysis described in resequencing and population analysis section. The nucleotide substitution model PMB+F+G4 was selected for *Ib*T-DNA1 and the model TVM+F was selected for *Ib*T-DNA2.

BEDTools genomecov (version v2.25.0) ^55^ was used to calculate the sequencing depth of different sites and recorded as bedgraph file. The INDELs (insertion and deletion variations) were extracted from vcf files and integrated in the bedgraph file. R package ggplot2 ^55^ was used to visualize the marked bedgraph files, which displayed the schematic diagram of the T-DNA structure and INDELs of T-DNA-positive accessions.

### Chloroplast genome assembly and phylogenetic analysis

The chloroplast genomes were assembled using GetOrganelle (version 1.7.5) ^65^, and chloroplast size ranges from 160,892 to 161,955 bp. The chloroplast genome sequences were aligned by MAFFT ^61^ by default, followed by revised align using MUSCLE ^66^ implemented in MEGA X ^63^. Gblocks (version 0.91b) ^67^ was used to remove poorly aligned positions (−b4=5 −b5=h), resulted in final 161,200 bp alignment length. The phylogeny was reconstructed using IQ-TREE (version 1.6.12) ^52^ with 1,000 ultrafast bootstrap replicates. The nucleotide substitution model (TVM+F+I) was selected by IQ-TREE. The chloroplast network was generated using the TCS Network method implemented in PopART (version 1.7) ^68^.

## Data availability

The raw DNA sequencing data are deposited in BIGD under accession number PRJCA004953.

## Code availability

The HPA pipeline and relevant instructions are available at the Github website (https://github.com/YanMengxiao/HPA). Other analysis command lines are given in the Supplementary Data file.

## Acknowledgements

This work was funded by the Ministry of Science and Technology of China (2019YFD1000703 to J.Y., 2019YFD1000704-2 to M.X.Y., and 2019YFD1000701-2 to W.J.F.), the Innovation Promotion Foundation of Sichuan Academy of Agricultural Sciences (2019LJRC017, 2019QYXK006 to M.L.), the National Natural Science Foundation of China (31771854 to H.X.W.), the Shanghai Municipal Afforestation & City Appearance and Environmental Sanitation Administration (G182402, G192413, G192414 and G202402 to J.Y.), the State Key Laboratory of Subtropical Silviculture (KF2019 to J.Y.), Youth Innovation Promotion Association CAS, and the Bureau of Science and Technology for Development CAS (KFJ-BRP-017-42 to J.Y.). The authors thank NARO genebank (Japan) for providing dried leaves of their materials, especially Dr. Masaru Tanaka and Dr. Hiroshi Nemoto.

## Author contributions

M.X.Y. involved in the conception of this study, developed HPA pipeline, conducted most of the data analysis and wrote the manuscript. M.L. provided plant materials, part of sequencing data. Y.Z.W. assembled chloroplast genomes and performed the T-DNA analysis. X.Y.W reconstructed the chloroplast network. M.H.M. updated the Ranbow software. D.G.Q. performed the PCR screening of most plants. W.J.F. performed the PCR screening and prepared the materials for RNA-seq. Y.Q.W. involved in assembly of chloroplast genomes of MDP. H.Z.N. performed the PCR screening of some samples. Z.Y.W. helped to access the plants and DNA samples. B.H. tracked back the collecting information of CIP accessions. R.J. provided the Ecuador accessions. J.F.K. and G.G. discussed and contributed to the early phase of the project. H.X.W. and R.B. revised the manuscript. R.B. suggested and contributed to the gene conversion analysis. M.V. discussed and contributed to the haplotyping analysis. J.Y. designed the study and contributed to the original concept of the project.

